# Development of an Engineered *Mycobacterium tuberculosis* Strain for a Safe and Effective Tuberculosis Human Challenge Model

**DOI:** 10.1101/2023.11.19.567569

**Authors:** Xin Wang, Hongwei Su, Joshua B. Wallach, Jeffrey C. Wagner, Benjamin Braunecker, Michelle Gardner, Kristine M. Guinn, Thais Klevorn, Kan Lin, Yue J. Liu, Yao Liu, Douaa Mugahid, Mark Rodgers, Jaimie Sixsmith, Shoko Wakabayashi, Junhao Zhu, Matthew Zimmerman, Véronique Dartois, JoAnne L. Flynn, Philana Ling Lin, Sabine Ehrt, Sarah M. Fortune, Eric J. Rubin, Dirk Schnappinger

## Abstract

Human challenge experiments could greatly accelerate the development of a tuberculosis (TB) vaccine. Human challenge for tuberculosis requires a strain that can both replicate in the host and be reliably cleared. To accomplish this, we designed *Mycobacterium tuberculosis* (Mtb) strains featuring up to three orthogonal kill switches, tightly regulated by exogenous tetracyclines and trimethoprim. The resultant strains displayed immunogenicity and antibiotic susceptibility similar to wild-type Mtb under permissive conditions. In the absence of supplementary exogenous compounds, the strains were rapidly killed in axenic culture, mice and nonhuman primates. Notably, the strain that contained three kill switches had an escape rate of less than 10^-10^ per genome per generation and displayed no relapse in a SCID mouse model. Collectively, these findings suggest that this engineered Mtb strain could be a safe and effective candidate for a human challenge model.

## Introduction

Tuberculosis (TB) stands as a leading global infectious threat, with its toll on lives and health exacerbated during the COVID-19 pandemic. Mathematical modeling underscores the urgency of developing novel TB vaccines to achieve the ambitious goals set by the "Global Plan to End TB" initiative by Stop TB Partnership^1^. The only licensed TB vaccine remains the attenuated *Mycobacterium bovis* strain known as Bacille Calmette-Guerin (BCG). Despite its efficacy against TB meningitis and miliary TB in infants^2^, BCG’s effectiveness against TB in adolescents and varies geographically^3^. An alternative vaccine candidate, M72/AS01^4^, exhibited protection rates against active TB no higher than 50% in adults with latent *Mycobacterium tuberculosis* infection, considerably lower than the 90% efficacy seen with polio and measles vaccines^5, 6^.

The challenges hindering TB vaccine development are two-fold: (1) the absence of inexpensive and predictive preclinical animal models and (2) the lack of validated immunological markers for protection. While progress has been made in understanding immune protection indicators^7, 8, 9, 10^, leveraging non-human primates as models, a <u>C</u>ontrolled <u>H</u>uman Infection <u>M</u>odel (CHIM), if it can be developed, would offer a valuable tool to assess vaccine efficacy and complement animal studies.

The CHIM, also referred to as the human challenge model, is a pivotal instrument in advancing vaccine development and gauging treatment efficacy. In this strategy, volunteers are deliberately infected with an infectious agent to evaluate the efficacy of vaccines or therapeutic candidates. Human challenge studies possess advantages not reproducible in natural infection investigations. By eliminating variables such as diverse genetic backgrounds of infectious agents, variable infectious doses, uncertain exposure periods, and patient co-morbidities, researchers can pinpoint protective host factors and immediate responses during infection. CHIM facilitates meticulous control over infection rates and timing, enabling comparisons among new vaccine candidates, modified regimens, and preventive, preemptive, or post-symptomatic treatments. Its success includes the development of various FDA-approved vaccines and therapies, such as the cholera vaccine Vaxchora^11^, the influenza therapeutic Oseltamivir^12^, the Vi-tetanus toxoid conjugate vaccine for *Salmonella typhi*^13^, and dosing schedules and adjuvant selections for malaria vaccines RTS,S and S/AS01^14, 15^. A recent BCG human challenge model conducted in South Africa introduced live BCG or purified protein derivative (PPD) into lung segments of volunteers through bronchoscopy, followed by immune profiling studies^16^. The observed adverse effects were mild, marking an important step toward establishing a CHIM in the realm of TB research.

A successful CHIM study must align with the ethical principle of "*primum non nocere*"^17, 18^. For TB CHIM, this requires engineering of an attenuated Mtb strain that adheres to preclinical safety standards. This challenge strain needs to be capable of replication in the naïve host to ensure that its eradication requires vaccine-boosted immune responses. Moreover, the challenge strain should permit only a limited window for replication to adhere to safety regulations, with the genetic circuit governing strain mortality demonstrating a low escape mutation rate (we set a stringent goal of <10^-12^ per genome per generation). To attain this objective, we designed and integrated three distinct kill switches employing orthogonal mechanisms into the Mtb strain H37Rv. The resultant triple-kill-switch (TKS) strain exhibited a growth rate and antibiotic susceptibility similar to the wild-type H37Rv under permissive conditions and was rapidly killed both *in vitro* and within mouse models under restrictive conditions. We determined that the escape rate for all three kill switches remained below our current limit of detection (<10^-10^ per genome per generation). Importantly, the TKS strain did not induce relapse in immune-compromised SCID mice. Based on our safety and efficacy assessments, we proposed the TKS strain as a promising candidate for a human challenge strain.

## Results

### Construction of tetracycline addicted dual lysin Mtb

We constructed two genetic switches that depend on a tetracycline (Tc), such as anhydrotetracycline (aTc) or doxycycline (doxy), to repress the lysin operons, L5L and D29L, of two mycobacteriophages, L5 and D29. L5L was repressed by a single-chain reverse TetR (revTetR)^19^ (Figure 1A). D29L was repressed by a modified TetPipOFF system^20^ in which a single-chain wild-type TetR represses PipR, which in turn controls activity of the D29L operon. To ensure independent regulation of the lysin operons, the two TetRs chosen were distinct in their operator specificity (revTetR binds to the operator mutant *tetO*_4C5G_ while TetR binds to wild-type *tetO*) and in their allosteric response to aTc/doxy (revTetR requires aTc/doxy to be active whereas TetR is inactivated by aTc/doxy). Single-chain repressors were used to prevent heterodimerization of the two TetRs^21^. The resulting “dual-lysin strain” is a derivative of Mtb H37Rv that carries both switches integrated in its genome.

**Figure 1.**
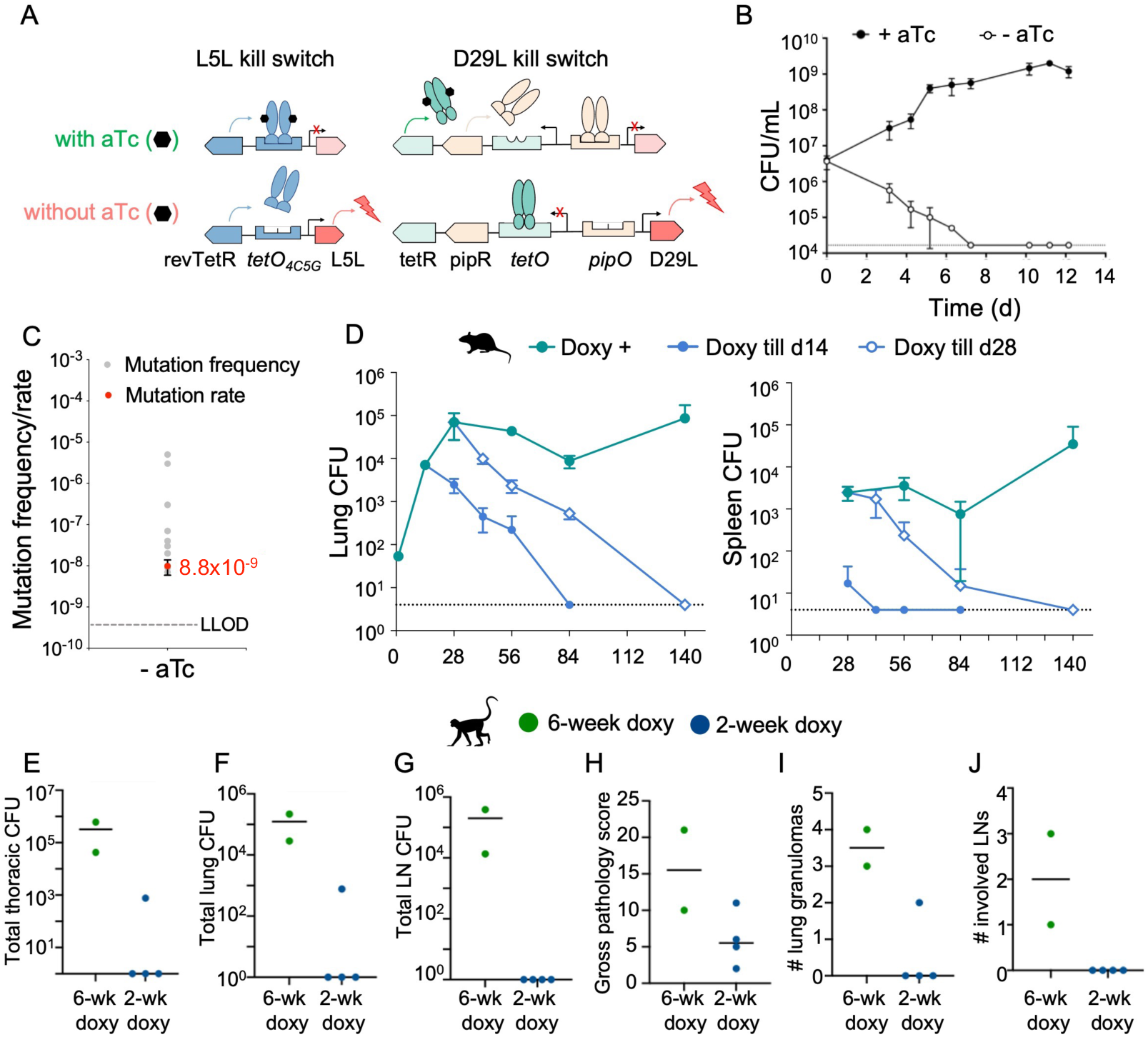
The dual-lysin kill switch is effectively bactericidal in *Mtb* both *in vitro* and *in vivo.* (**A**) Regulatory scheme of aTc-inducible lysin expression. (**B**) The kill curve of dual-lysin *Mtb* strain *in vitro.* (**C**) The escape mutation rate of dual-lysin *Mtb* strain in the absence of aTc. (**D**) The clearance kinetics of *Mtb* dual-lysin strain in C57BL/6 mice with doxy depleted at day 14 or day 28. Error bars represent standard deviation in each panel. (**E** to **J**) The NHP infection outcome of the dual-lysin *Mtb* strain supplemented with doxy for 2 weeks or 6 weeks. The outcome is judged by (**E**) total thoracic CFU, (**F**) lung CFU, (**G**) lymph node (LN) CFU, (**H**) gross pathology score, (**I**) number of lung granulomas at necropsy and (**J**) number of involved lymph nodes. Each symbol represents a macaque and the line represents median.

In axenic culture, the dual-lysin strain exhibited growth in the presence of aTc, while in its absence, it displayed potent bactericidal effects, with a kinetic rate of a ∼1-log reduction in CFU every 3 days (Figure 1B). Through fluctuation assays, we determined the *in vitro* escape rate as 8.8 x 10^-9^ per genome per generation (Figure 1C). In a C57BL/6 murine model, the dual-lysin strain effectively established an infection and persisted as long as doxy was administered. Following doxy withdrawal, the strain was cleared from both lungs and spleen with killing kinetics of approximately 0.50 log/week and 0.35 log/week, respectively (Figure 1D). In non-human primates (NHP, here Mauritian cynomolgus macaques) using the Mtb Erdman strain with the dual lysin constructs, a 6-week doxy regimen resulted in infection reaching up to 10^6^ CFU in the thorax, lungs, and spleen. Shortening the doxy supplementation to 2 weeks led to sterilization of all assessed organs, except for a solitary instance of 10^3^ CFU in the thorax and lungs of one animal (Figures 1E to 1H). Notably, the presence of viable colonies in the one NHP under the 2-week doxy treatment was not attributed to escape mutations, as we verified that bacterial growth was still regulated by aTc in axenic culture. Lung granulomas were only found in 1 of the 4 macaques on the 2-week doxy regimen, while both macaques on doxy for the duration of the study had visible lung granulomas (Figure 1I). Similarly, none of the 2-week doxy regimen macaques had thoracic lymph nodes with signs of disease while both 6-week doxy regimen macaques had involved lymph nodes (Figure 1J).

Collectively, these findings underscore the stringent regulation of the dual-lysin strain’s viability by aTc, both *in vitro* and *in vivo*. However, the kinetics of bacterial elimination were not as swift as desired, and the escape rate surpassed our benchmark, thereby raising safety concerns regarding the strain’s suitability for human challenge studies. To achieve the target safety standards, our focus shifted towards developing a third kill switch.

### Engineering degron-mediated Mtb proteolysis via ClpC1-P1P2 protease controlled by trimethoprim

To design a killing mechanism orthogonal to lysin expression, we sought to leverage degradation of essential and vulnerable Mtb proteins. Prior research has described a degron engineered to mediate protein degradation in mammalian central nervous system^22^. This degron was characterized by a mutated *E. coli* dihydrofolate reductase (DHFR) that forms an unstable protein conformation, consequently driving protein degradation. In the context of eukaryotic system, proteolysis was hindered by trimethoprim (TMP), an ineffective antibiotic against Mtb^23^, which binds to the degron (ddTMP), to stabilize its unfolded conformation. To investigate the applicability of the ddTMP degron in mycobacteria, we fused a red fluorescent protein (mCherry in *Mycobacterium smegmatis* (Msm) or mScarlet in Mtb, both designated RFP) with ddTMP at both the C- and N-termini and evaluated the TMP-dependent stability of the resultant RFP construct in Msm. In the absence of TMP, RFP was susceptible to proteolysis when ddTMP was fused at either the N- or C-terminus but the presence of TMP prevented degradation of RFP (Figures 2A, 2B and 2C). In Mtb, the ddTMP-RFP was similarly stabilized in a TMP-dependent manner (Figure 2D).

**Figure 2.**
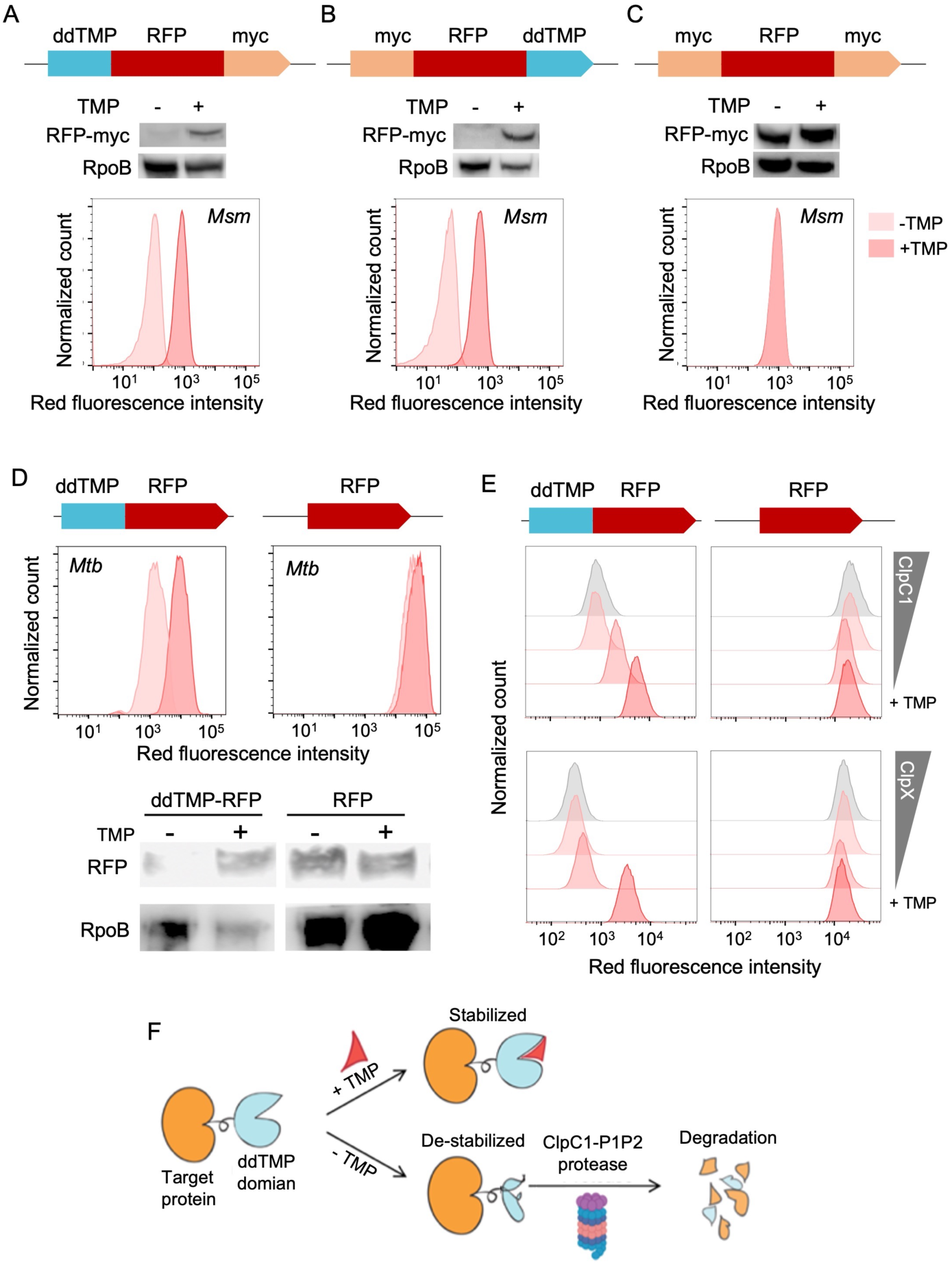
Mycobacteria proteins fused with ddTMP tag are subjected to trimethoprim (TMP)-dependent degradation via ClpC1-ClpP1P2 protease. (**A** to **C**) mCherry fused with ddTMP at N-terminus (**A**), C-terminus (**B**) or null (**C**). The ddTMP at either N-terminus or C-terminus manifests TMP-dependent degradation in *M. smegmatis.* (**D**) ddTMP-mScarlet shows a TMP-dependent degradation in *M. tuberculosis.* (**E**) Flow cytometry with *clpC1* or *clpX* knockout *M. smegmatis* indicates that the TMP-dependent degradation is mediated via protease ClpC1-ClpP1P2. The bottom row shows RFP steady state measured in mid-log culture supplied with TMP. *ClpX* or *clpC1* were knocked down via a gradient of aTc in the absence of TMP. (**F**) shows the ClpC1-dependent degradation mechanism of the ddTMP tag in mycobacteria.

To decipher whether the ddTMP-mediated proteolysis was dependent on ClpC1 or ClpX acting as the ATPase with the ClpP protease, we analyzed the degradation kinetics of ddTMP-RFP in aTc-induced *clpC1* or *clpX* knockdown Msm strains. Our data indicated that depleting ClpC1 led to the accumulation of ddTMP-RFP. Conversely, the abundance of ddTMP-RFP was unaffected by the depletion of ClpX (Figure 2E). These data indicate that ddTMP-tagged proteins undergo degradation primarily through ClpC1 as the cognate ATPase with the Clp protease system in Msm (Figure 2F).

### Trimethoprim-dependent Mtb NadE degradation is bactericidal *in vitro*

We sought to integrate the ddTMP degron into Mtb to engineer a kill-switch strain. We fused the ddTMP degron to the C-terminus of the essential Mtb protein NadE at its native chromosomal locus (Figure 3A). NadE is essential for catalyzing the final step in NAD^+^ *de novo* biosynthesis. NadE depletion has been shown to induce rapid killing during both replication and persistence^19^ as a result of the reduction of both NADPH and NADH pools^24^. We assessed the kinetics of NadE degradation using a reporter construct where NadE was fused with mScarlet-ddTMP at the C-terminus. Absent external TMP, we observed exponential degradation of NadE-mScarlet-ddTMP, with a half-life of ∼1 hour (Figure 3B). The minimum concentration of TMP required to support the growth of the NadE-ddTMP strain was 3 ng/mL in axenic culture. At TMP concentrations higher than 30 ng/mL, the NadE-ddTMP strain displayed no apparent growth defect compared to the wild-type H37Rv strain (Figure 3C). In the absence of TMP supplementation, the degradation of NadE had a bactericidal effect, evident in a 2-log reduction in CFU within a 12-day period *in vitro* (Figure 3D).

**Figure 3.**
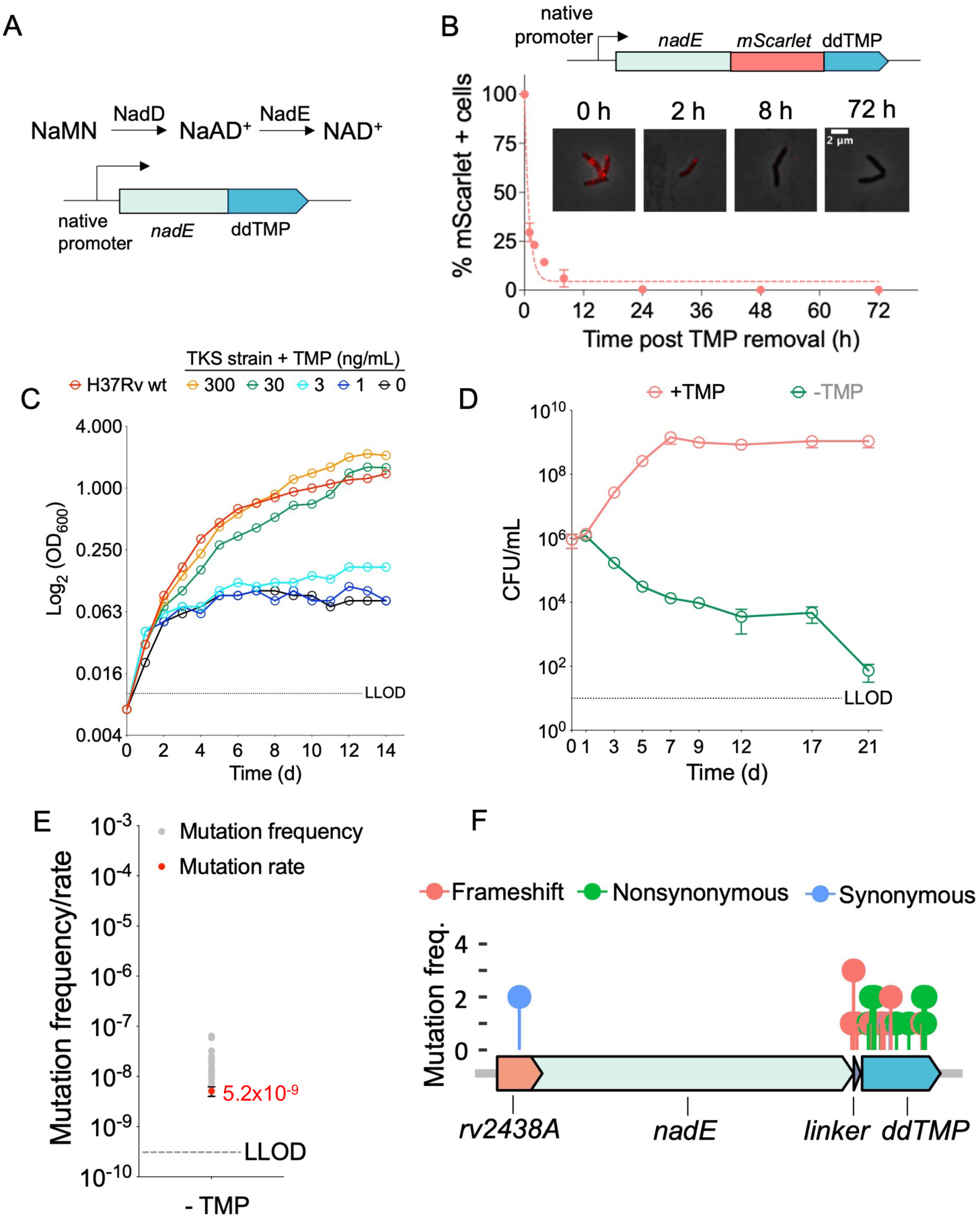
TMP-dependent NadE degradation leads to *Mtb* death. (**A**) The genetic construct of tagging *Mtb* gene *nadE* with C-terminus ddTMP. (**B**) The degradation kinetics of NadE-mScarlet-ddTMP were measured by quantifying mScarlet intensity per cell via microscope. The degradation curve is fitted into a one-phase exponential decay model. (**C**) The growth curve of *Mtb nadE-ddTMP* strain with various TMP concentrations in 7H9 media. (**D**) The kill curve of *Mtb nadE-ddTMP* strain in the absence of TMP. (**E**) The mutation rate of *Mtb nadE-ddTMP* strain and (**F**) its escape mutation density plot are determined by whole-genome sequencing. Error bars represent standard deviation in each panel.

Since we observed TMP-regulated growth and killing, we assessed the escape mutation rate and characterized the mutations leading to escape. Through fluctuation assays, we determined the escape rate of the NadE-ddTMP kill switch to be ∼5.1 x10^-9^ per genome per generation (Figure 3E). Subsequent whole-genome sequencing of 20 escape mutants of NadE-ddTMP revealed that most resistance mutations were frameshift or nonsynonymous mutations at the *nadE* C-terminus or within the ddTMP degron region (Figure 3F and Supplemental Table 1). This pattern indicated that escape likely occurred due to the functional ablation of the ddTMP degron. We also identified a synonymous mutation in the *nadE* upstream region, which likely reflects a promoter mutation inducing higher *nadE* expression to counteract proteolysis stress (Figure 3F, and Supplemental Table 1). These escape mutations indicate that the Mtb killing effect was indeed attributed to NadE degradation, devoid of off-target consequences.

### Induction of the combined triple-kill-switch sterilizes Mtb cultures *in vitro*

A successful Mtb human challenge strain necessitates growth kinetics resembling the wild-type strain under permissive conditions, while ensuring rapid, low-escape-rate killing under restrictive conditions. To achieve this, we combined the two lysin switches with NadE-ddTMP, yielding a triple-kill-switch (TKS) Mtb H37Rv strain. The TKS strain’s growth kinetics closely mirrored those of wild-type H37Rv when cultivated under permissive conditions with aTc and TMP supplementation (Figure 4A). Under permissive conditions, disparities with small effect size (Cohen’s d < 0.200) in cellular morphology were observed between the TKS and H37Rv strains (Figure 4B). Furthermore, susceptibility to first-line and second-line antimycobacterial agents such as isoniazid, rifampin, ethambutol, and moxifloxacin remained indistinguishable between the TKS strain and H37Rv (Figures 4C to 4F). However, in the absence of aTc and TMP, the TKS strain exhibited rapid killing as reflected by an ∼3-log decline in CFU per week. In axenic culture, complete sterilization was achieved within 11 days after aTc and TMP removal (Figure 4G). Depleting either TMP or aTc individually did not yield such rapid and complete sterilization, implying a synergistic bacterial-killing effect between the phage-lysin and ddTMP degron kill switches.

**Figure 4.**
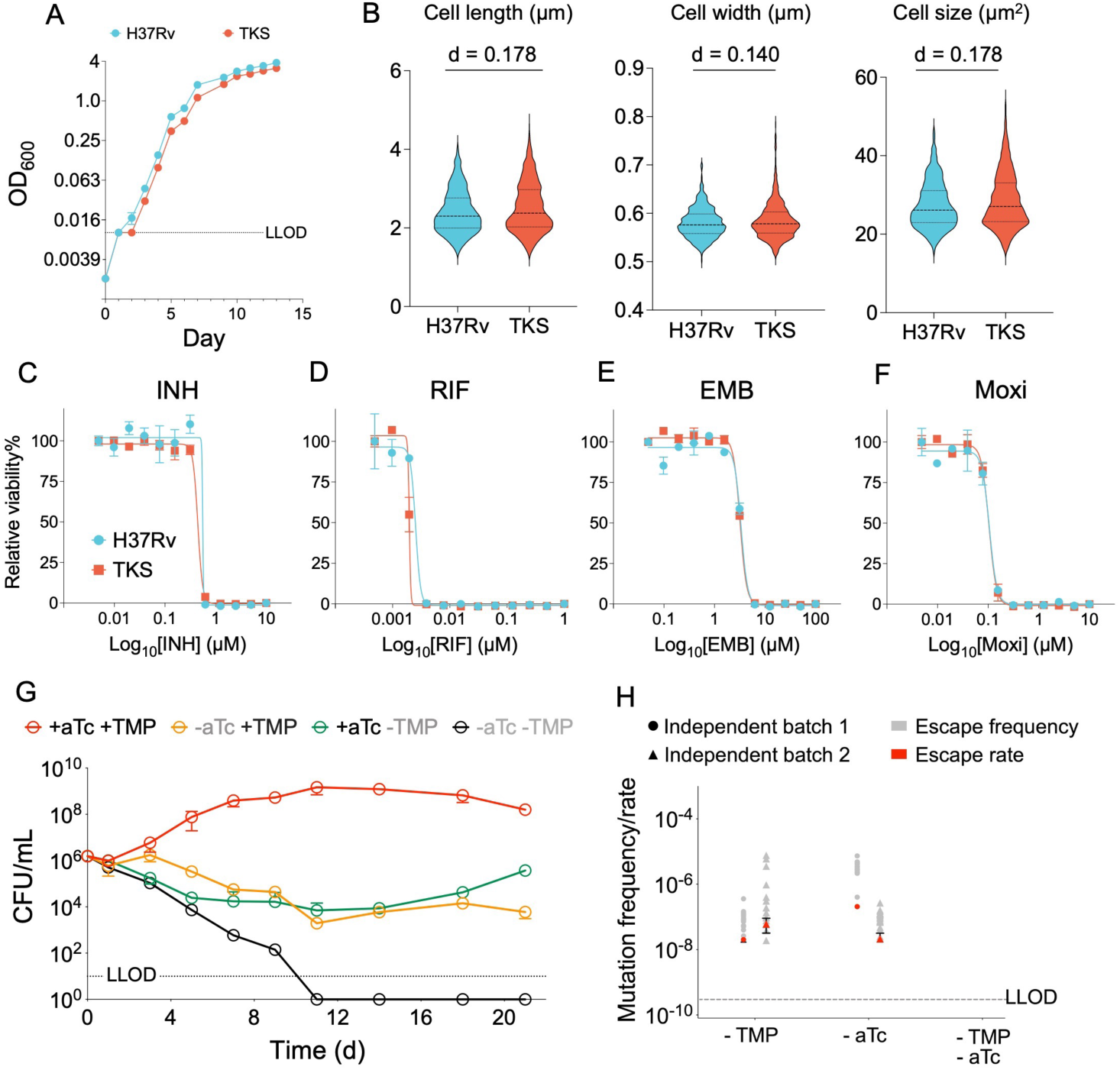
The triple-kill-switch (TKS) strain manifests normal fitness at permission condition yet rapid killing and low escape rate at restrictive condition. **(A)** The growth curve of TKS and H37Rv strains at permissive condition. The dash line shows the lower limit of detection as OD 0.001. **(B)** The quantified bacterial morphology of strains H37Rv (n=1068) and TKS (n=1614). Cohen’s d was calculated to show effect size between H37Rv and TKS strains. **(C)** to **(F)** The MIC of H37Rv and TKS strains against isoniazid (INH), rifampin (RIF), ethambutol (EMB) and moxifloxacin (Moxi) at permissive condition. **(G)** The *in vitro* kill curve of TKS strain in the absence of aTc and/or TMP. The dash line shows the lower limit of detection as 10 CFU/mL. **(H)** Escape rate of TKS strain in the absence of aTc or/and TMP from two independent experiments. The dash line shows the lower limit of detection as 3.8x10^-10^. Error bars represent standard deviation in each panel.

The design of the TKS strain enabled us to measure the escape rates of each system independently. We determined escape rates with two independent batches of bacteria culture using fluctuation analysis. In the presence of aTC, we found the escape rate for the ddTMP degron ranging from 2.0x10^-8^ to 6.0x10^-8^ per genome per generation. In the presence of TMP, we found the dual-lysin escape rate ranging from 2.1x10^-8^ to 2.1x10^-7^ per genome per generation (Figure 4H). Whole genome sequencing indicated that escape mutations in the ddTMP degron system resulted from the truncation of the ddTMP degron tag, while escape mutations associated with the dual-lysin kill switches arose from disruptions and deletions within the lysin operons (Supplemental Table 2). Importantly, the escape rate fell below the lower limit of detection within our current experimental setup under the stringent restrictive conditions, encompassing depletion of both aTc and TMP (< 3.8x10^-10^ per genome per generation). Since escape mutation mechanisms are independent, the theoretical escape rate should be <10^-15^.

### The triple-kill-switch strain is rapidly cleared in mice without discernible relapse

To test the TKS strain virulence in permissive conditions and its clearance in restrictive conditions, we infected C57BL/6 mice with the TKS strain via the aerosol route. PK/PD studies on doxy and TMP suggested that dietary supply or removal of doxy and TMP promptly modulated their levels in mouse tissues and plasma, either above or below the threshold necessary for the TKS strain survival^25^ (Supplemental Table 3). Therefore, we fed mice chow supplemented with TMP and doxy for 28 days and then switched to regular chow until day 140 (Figure 5A). The TKS strain established infection under permissive conditions, resulting in pulmonary CFU ranging from 10^3^ to 10^4^ by day 28. The pulmonary burden under permissive conditions reached a plateau of 10^5^ CFU (Figure 5B). Dissemination to the spleen was observed at ∼day 28, as evident from 2 out of 5 mice with a spleen bacterial burden of ∼100 CFU. The spleen CFU stabilized at approximately 10^3^ to 10^4^ CFU after day 56 (Figure 5C). Upon discontinuing TMP and doxy supplementation at day 28, the pulmonary bacterial burden fell at a rate of ∼0.75 log CFU reduction per week.

**Figure 5.**
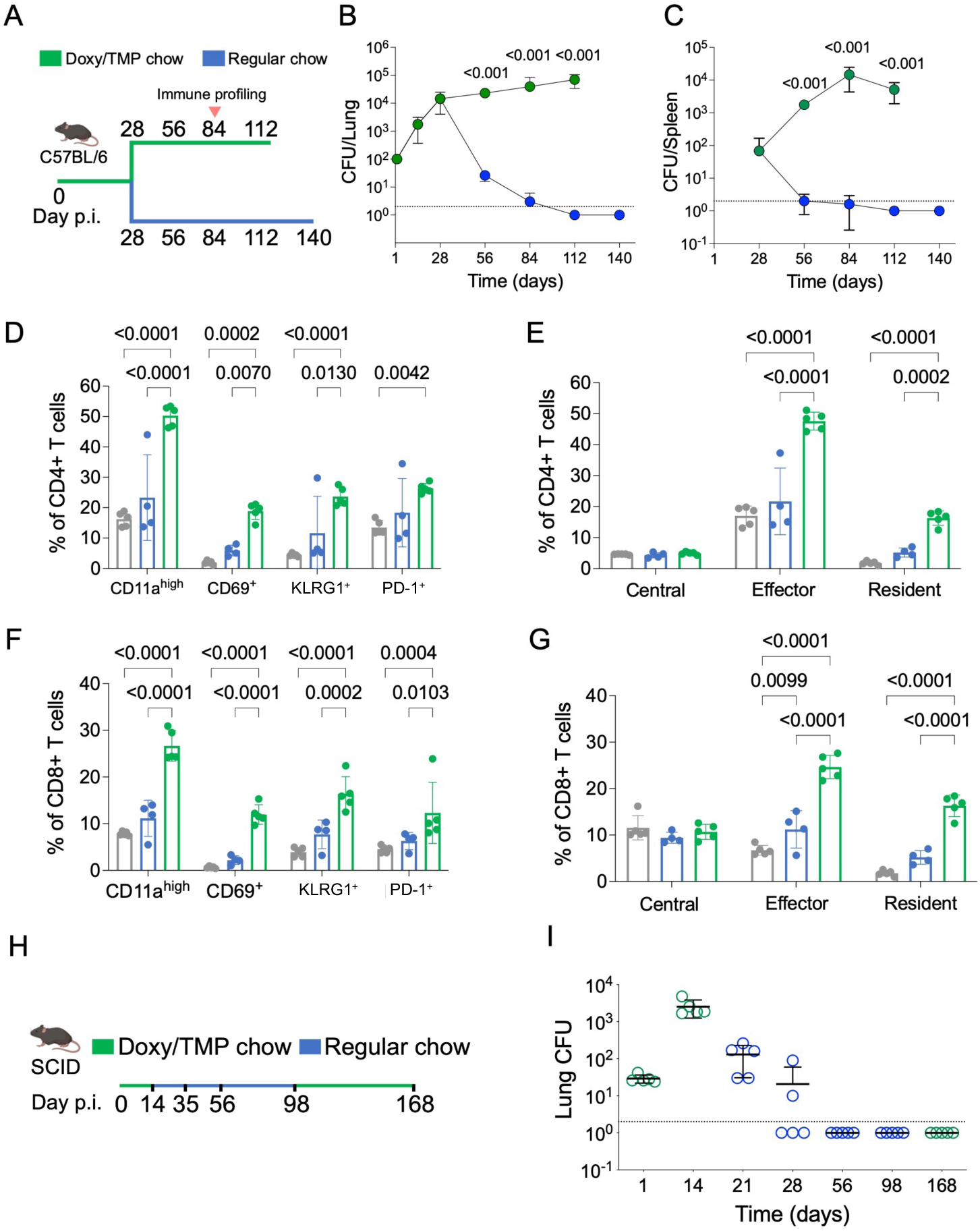
The combo kill switch strain is rapidly cleared in mouse without discernible relapse. (**A**) The scheme of clearance study in C57BL/6 mice. Mice were kept on doxy/TMP chow 3 days before infection till day 28 post-infection. Lung and spleen CFU were enumerated at indicated time point. (**B** and **C**) Lung and spleen CFU of the C57BL/6 mouse experiment. P-values were calculated via Student’s t-test. (**D**) The proportion of CD4 T cells with activation markers at day 84. (**E**) The proportion of CD4 memory T cells at day 84. (**F**) The proportion of CD8 T cells with activation markers at day84. (**G**) The proportion of CD8 memory T cells at day84. P-values in (D) to (G) were calculated by Student’s t-test with multiple comparison corrected with Tukey method. (**H**) The scheme of relapse study in SCID mice. Mice were kept on doxy/TMP chow 3 days before infection till day 14 post-infection. Lung CFUs were enumerated at indicated time points. (**I**) The kinetics of lung CFU enumerated in the SCID relapse experiment.

Extrapolation suggested complete pulmonary clearance by day 96, with no detectable live bacteria at days 112 and 140 under restrictive conditions (Figure 5B). Only 1 out of 5 mice had detectable bacteria in the spleen on days 56 and 84, and all mice cleared splenic infection on days 112 and 140 (Figure 5C).

To investigate immune responses, we examined pulmonary CD4^+^ and CD8^+^ T cell profiles on day 84 in mice with or without doxy/TMP supplementation. TKS infection under permissive conditions prompted infiltration of activated T cells (CD11a^high^, CD69^+^, KLRG1^+^, and PD-1^+^) (Figures 5D, 5F and Supplemental Figure 1), along with the recruitment of effector and resident memory T cells (Figures 5G, 5E and Supplement figure 1). Although this response was attenuated under restrictive conditions, it remained distinguishable from uninfected mice.

To assess the potential for relapse under stringent conditions, we utilized the immunocompromised SCID mouse model (Figure 5H). We administered TMP and doxy supplemented chow from days 1 to 14 and days 98 to 168. The TKS strain established infection with 5x10^3^ pulmonary CFU by day 14. Upon depleting TMP and doxy, bacteria cleared at a rate of ∼1.5 log CFU per week. There was no detectable bacterial growth from lung homogenates at days 56 and 98. After return to doxy and TMP supplementation for 10 weeks, no viable bacteria were detected at day 168 in the lungs (Figure 5I) or spleen (Supplemental Figure 2). Thus, the TKS strain did not reemerge under permissive conditions, even in the absence of adaptive immunity, once it had been eliminated by kill switch induction.

In conclusion, we observed tight regulation of TKS strain growth and death through doxy/TMP supplementation, with discontinuation leading to rapid clearance and attenuation of the immunopathological response. Importantly, no relapse occurred even in the absence of an intact immune system.

## Discussion

The burgeoning interest in Controlled Human Infection Model (CHIM) studies, aimed at expediting TB vaccine development, has emphasized the critical need for an engineered Mtb challenge strain to effectively address safety concerns. When assessing vaccine efficacy, the challenge strain must demonstrate consistent replication to ensure robust immunogenicity, while also enabling rapid killing at the conclusion of the study. This rationale led us to design a challenge strain reliant on exogenous molecules, triggering bacterial killing upon the removal of small molecule supplementation. To meet these goals, we have engineered an Mtb strain equipped with three distinct kill switches. This engineered strain exhibits growth kinetics and antibiotic susceptibility akin to the wild-type Mtb in the presence of doxy and TMP. In the absence of doxy and TMP, it is rapidly eradicated *in vivo*.

Safety is the foremost concern for the TB human challenge strain. The most noteworthy attribute of the TKS strain is its robust killing in the absence of doxy and TMP supplementation. The removal of doxycycline prompts the expression of lysins L5 and D29, culminating in cell wall damage and bacterial lysis. The absence of TMP triggers ClpC1-mediated degradation of NadE, a crucial metabolic enzyme in the final step of NAD^+^ biosynthesis. The amalgamation of these three orthogonal killing mechanisms yields a low escape rate and rapid elimination kinetics. Because single mutations could arise during passage, we set a rigorous goal for the escape rate as <10^-12^. The calculated rate derived from experimental measurements exceeds that goal.

A second safety goal is to attain rapid clearance of bacteria after withdrawal of antibiotics. We saw rapid clearance rates in mice regardless of their immune status. Importantly, the TKS strain did not cause relapse disease in immunocompromised SCID mice, even after ten weeks of TMP and doxy reintroduction. Collectively, these growth and killing attributes derived from the kill switches suggest that this strain could be safely used for TB human challenge.

Thus far, no animal model reliably predicts vaccine efficacy in humans. Although mice are widely used in TB immunology studies, they fail to replicate human TB disease due to cross-species differences such as distinct macrophage responses, the absence of pulmonary granuloma structures, and differing antigen presentation repertoires. These disparities impede TB vaccine development grounded solely in mouse studies. Non-human primate (NHP) studies model human TB better than mice, though, again, we do not yet know their predictive power. A human challenge model may allow more rapid identification of human immune correlates in TB infection and protection, facilitating vaccine efficacy testing and validating correlates identified in previous NHP and mouse studies. A recent human challenge study in the UK, involving the bronchial inoculation of BCG or PPD in humans, revealed novel insights distinct from prior *in vitro* studies^16^. However, a limitation to the use of BCG as a CHIM strain is the lack of expression of certain antigens, particularly those encoded in the RD1 region, some of which are included in vaccine platforms. Thus, an Mtb challenge strain would be more appropriate in assessing vaccine effectiveness. Future TB CHIM work using the attenuated Mtb TKS strain could illuminate host-pathogen interaction more relevant to Mtb infection than the BCG and PPD model.

Our study has limitations. Firstly, we have not evaluated the safety of the TKS strain in other mammalian models, such as NHPs. Assessing kill kinetics and relapse rates in NHPs will help validate safety for human studies. It will also be important to determine a “safe” bacterial burden that minimizes the risk of pathologic injury to participants. Secondly, alternative challenge routes, apart from aerosol, could be explored for the TKS strain to assess its ability to detect a vaccinal effect. Considering the distinct tissue pharmacokinetics and pharmacodynamics of doxy and TMP, it will be important to benchmark growth and killing kinetics in multiple tissues for the TKS strain. Thirdly, a sensitive and non-invasive detection method will be important to track strain clearance *in vivo*. Ideally, the detection approach should be non-invasive to minimize costs and efforts for clinical cohorts and volunteers.

In conclusion, we have successfully engineered an Mtb triple kill-switch strain with immunogenicity comparable to the wild-type strain under permissive conditions and displaying rapid and complete sterilization without relapse under restrictive conditions. We propose the TKS strain as a suitable challenge strain for a TB controlled human infection model, advancing TB vaccine efficacy studies and shedding light on human immune responses to Mtb infection.

## Methods

### Strains, media, and culture conditions

All Mtb strains are H37Rv derivatives, except the dual-lysin strain used for macaque studies which was constructed in the Erdman background. All Msm strains are mc^2^155 derivatives. The wild-type strains H37Rv or mc^2^155 were grown in Middlebrook 7H9 broth or 7H10 agar suppled with 0.5% glycerol, 0.05% Tween-80 and 1x OADC (oleic acid-albumin-dextrose-catalase, Middlebrook 212351) supplement (Mtb) or 1x ADC (albumin-dextrose-catalase) supplement (Msm). For the permissive condition of the dual-lysin strain, the media was supplemented with aTc (0.5-1.0 µg/mL), kanamycin (20 µg/mL) and zeocin (25 µg/mL). For the permissive condition of the NadE-ddTMP strain, the media was supplemented with TMP (50 µg/mL) and hygromycin (50 µg/mL). For the TKS strain, the media was supplemented with aTc (0.5-1.0 μg/mL), TMP (50 µg/mL), kanamycin (20 µg/mL), hygromycin (50 µg/mL) and zeocin (25 µg/mL).

### Generation of strains

To create the dual-lysin kill switch Mtb strain, H37Rv or Erdman was transformed with plasmids pGMCK3-TSC10M-TsynE-pipR-SDn-PptR-D29L and pGMCgZni-TSC38S38-P749-10C-L5L, incorporating the D29-lysin and L5-lysin into Mtb L5 and Giles sites, respectively. ATc was supplemented at 0.5-1.0 µg/mL.

To create the NadE-ddTMP Mtb strain, H37Rv with plasmid pKM461 for ORBIT recombineering^26^ was transformed with an oligo (Supplementary Table 4) and a payload plasmid pJW461. The oligo contains an *attP* site flanked by 70 nucleotides at 5’- and 3’- end respectively homologous to the insertion site. The pKM461 contains an *attB* site and ddTMP sequence. These constructs resulted in labeling the ddTMP tag at C-terminus of NadE mediated via ORBIT as described previously^26^. The NadE-ddTMP strain was maintained with TMP supplemented at 50 µg/mL.

To create the triple-kill-switch (TKS) strain, the NadE-ddTMP strain was transformed with plasmids pGMCK3-TSC10M-TsynE-pipR-SDn-PptR-D29L and pGMCgZni-TSC38S38-P749-10C-L5L to incorporate the D29-lysin and L5-lysin into the Mtb genome with aTc supplemented at 0.5 µg/mL.

### Fluctuation analysis

Fluctuation analysis was performed as previously reported^27^. Briefly, the Mtb strain was inoculated at permissive condition (0.5-1.0 µg/mL aTc and 50 µg/mL TMP) in 7H9 media supplemented with OADC in the presence of antibiotics (20 µg/mL kanamycin, 25 µg/mL zeocin and 50 µg/mL hygromycin). After reaching OD=1.0, the culture was diluted to multiple 4-mL aliquots with 10,000 bacteria. The diluted culture was grown for 11-14 days in 7H9+OADC media in permissive conditions in the presence of antibiotics. Once OD was at 1.0, bacteria were washed for three times and resuspended in 400 µL 7H9+OADC without aTc or TMP. Four aliquots of bacteria were streaked onto 7H10+OADC plates supplemented with 0.5-1.0 µg/mL aTc and 50 µg/mL TMP for bacteria count, and the rest aliquots were spread onto 7H10+OADC plates with either aTc or TMP, or neither supplement. According to Ma, Sarkar, Sandri (MSS) method^28, 29^, the estimated number of mutations per culture (*m*) was inferred by number of mutant (*r*) colonies observed on plates. The escape rate was calculated by dividing *m* by *N_t_*, the number of cells plated for each culture. The Mann-Whitney *U* test was used to statistically compare escape rates between two groups. The lowest detection limit was calculated based on an assumption that only one colony could be observed in all 20 independent cultures.

### Mtb flow cytometry

Mtb cells were fixed in 2% paraformaldehyde overnight and removed from the BSL-3 facility. Fixed bacilli were quenched with 200 mM Tris-HCl (pH 7.5) for 5 minutes at room temperature and resuspended in PBST buffer (1x PBS with 0.1% Triton X-100). To suppress signals from noise or cell debris, two event triggers (thresholds) on forward scatter peak height (FSC-H >1.5) and side scatter area (SSC-A > 1.0) were used upon recording. To remove cellular aggregation, stringent gate settings were manually defined via FlowJo v. 10.8 to exclude events with strongly correlated forward scatter area (FSC-A) and SSC-A measures (large and compact particles), as well as events with disproportional FSC-A and FSC-H measures (morphological outliers). After event filtration, the log_10_-transformed red fluorescence intensity peak height (denoted TdTomato-A) was used to represent the abundance of intracellular red fluorescence protein.

### Microscope imaging and analysis

Mtb cultures were fixed with 2% paraformaldehyde (PFA) for 1 hour. PFA quenching and cell disaggregation were achieved by washing the fixed bacilli with a customized solution containing 200mM Tris-HCl (pH = 7.5), 1% (w/v) Triton X-100, 0.67% (v/v) xylenes and 0.33% (v/v) heptane^30^. Cells were then washed once with PBS-Triton (0.1% v/v) and seeded onto molded 1.8% agarose in phosphate buffered saline (PBS) for imaging. Phase contrast and fluorescence micrographs were collected via a Plan Apo 100× 1.45 NA objective using a Nikon Ti-E inverted, widefield microscope equipped with a Nikon Perfect Focus system with a Piezo Z drive motor, Andor Zyla sCMOS camera, and NIS Elements (v4.5). mScarlet signal was acquired using a 6-channel Spectra X LED light source and the Sedat Quad filter set. The excitation (Ex.) and emission (Em.) filters were Ex. 550/15nm and Em. 595/25nm, respectively. Semi-automated imaging was carried out using a customized Nikon JOBS script to locate imaging fields of interest, 9-12 images were taken for each sample. Cell segmentation and quantification was performed using our previously published Python pipeline, MOMIA^31^. We applied R function `lsr::cohensD()` to calculate effect size between H37Rv and TKS strains.

### Kill curve assay

Mid-log phase Mtb cultures (OD=0.3, 10^8^ CFU/mL) grown in permissive conditions were washed three times with 7H9+OADC without aTc or TMP, followed by dilution to 10^6^ CFU/mL (OD = 0.003). The diluted cultures were supplemented with aTc or TMP or both, and bacterial CFU was enumerated by plating serial dilutions on 7H10+OADC plates at multiple time points.

### Mtb genomic DNA extraction

Mtb culture at OD 1.0 was pelleted in buffer TE-NaCl followed by delipidation with chloroform:methanol (2:1). The delipidated bacteria were reconstituted in TE-NaCl followed by lysozyme (100 µg/mL) digestion at 37°C overnight and proteinase K (100 µg/mL) digestion in 0.01% SDS at 50°C for 2 hours. The digested lysate was treated with equal volume of phenol:choloroform (1:1, pH=8) followed by centrifugation, and the aqueous phase was treated with ½ volume of chloroform. After centrifugation, the gDNA in aqueous phase was precipitated with 1/10 volume of 3 M sodium acetate (pH=5.2) and one volume of isopropanol at -20°C overnight. The gDNA pellet was washed with 70% ethanol twice and dissolved in nuclease-free water (Invitrogen). The gDNA concentration was measured by Qubit fluorometer.

### Mtb whole genome sequencing and SNP calling

The gDNA library was prepared with Nextera XT DNA Library Preparation Kit (Illumina, FC-131-1096) and Nextera XT Index Kit v2 (Illumina, FC-131-2001). The pooled library was sequenced on MiSeq platform (Illumina) with paired-end strategy. Reads were trimmed by sickle (version 1.33)^32^ to preserve reads with a Phred base above 20 and read length longer than 30, followed by mapping against the H37Rv reference genome (ASM19595v2) using bwa^33^. The duplicated reads were then removed by SAMtools^34^. We used VarScan (version 2.3.9)^35^ to call SNP variants with the parameters: `--min-coverage 3 --min-reads2 2 --min-avg-qual 20 --min-var-freq 0.01 --min-freq-for-hom 0.9 --p-value 99e-02 --strand-filter 0`.

### Minimal inhibitory concentration

MIC values of drugs were determined following the microplate-based Alamar Blue assay (MABA) method as previously described^36^. Two-fold serial dilutions of drug were prepared in sterile polystyrene 96-well round-bottom plates (CLS3795, Corning, NY) with Middlebrook 7H9+OADC media, 100 µL per well. Mtb strains were 1:1000 diluted in 7H9+OADC medium at mid-logarithmic stage of growth (OD = 0.4). 50 µL of diluted bacterial suspensions were inoculated into each well, followed by incubation at 37°C for 7 days. 20 µL alamarBlue reagent (Invitrogen, Frederick, MD), freshly mixed with 12.5 µL 20% Tween 80, was added into each well, followed by 24-h incubation at 37°C. Absorbance was read at 570 nm, with reference wavelength 600 nm, using a microplate reader. The MIC endpoint was defined as the lowest concentration of the test agent that produced at least 90% reduction in absorbance compared with that of the drug-free control.

### Mouse experiments

#### Clearance study

C57BL/6J mice (The Jackson Laboratory, 000664) were aerosol infected with the triple-kill-switch (TKS) strain with 50-100 CFU. We used chow supplemented with 2000 ppm doxycycline and 1600 ppm TMP (Research Diets, New Brunswick, NJ) as permissive chow. The mice were fed with permissive chow starting 3 days before infection through 28 days post-infection. After day 28, mice on in the permissive arm were fed with permissive chow through day 112, whereas mice in the restrictive arm were fed with regular chow through day 140. Bacterial burden was determined by plating lung and spleen homogenates at day 1, 28, 56, 84, 112 and 140 post-infection.

#### Lung cell isolation

Murine right lung lobes were harvested into 2mL cold PBS in 5mL conical tubes kept in a cooling rack. Once all lungs were harvested, right lung lobes were transferred into gentleMACS C tubes (Miltenyi Biotec) containing 2.5mL pre-warmed digestion buffer consisting of filter-sterilized PBS (with calcium and magnesium) 0.5% bovine serum albumin (BSA) with freshly added DNase I (Roche, 10104159001) at 100μg/mL and Collagenase IV (Worthington, LS004186) at 150 units/mL. Lungs were dissociated in a gentleMACS Dissociator (Miltenyi Biotec), followed by incubation at 37°C for 45min. The digested tissue was dissociated again and passed through a 70µm cell strainer into a 50mL conical tube. The strainer was rinsed with 2mL of PBS 0.5% BSA. Red blood cells were lysed with eBioscience 1X RBC Lysis Buffer (Thermo Scientific 00-4333-57). Lung cells were washed in PBS, resuspended in RPMI complete medium (RPMI-1640 supplemented with 10% FBS, 2mM GlutaMax, 10mM HEPES, 0.05mM 2-mercaptoethanol and 1% Pen/Strep) and enumerated using the automated cell counter Countess II (ThermoFisher Scientific).

#### Flow cytometry of lung cells

Lung cell suspensions were washed with PBS twice, stained for viability with the live/dead dye Zombie UV (Biolegend, 423107) for 10 min at 4°C, washed with Cell Staining Buffer (BioLegend 420201) and incubated with Fc block (purified anti-mouse CD16/CD32 antibody, Biolegend 101302) at 1:200 in Cell Staining Buffer for 10 min at 4°C. After one wash in Cell Staining Buffer, cells were incubated with fluorochrome-conjugated monoclonal antibodies (mAbs) diluted at 1:200 into a 1:3 solution of Brilliant Staining Buffer (BD 563794): Cell Staining Buffer, for 45 min at 4°C. Cells were then washed twice in Cell Staining Buffer and fixed with Fixation Buffer (Biolegend 420801) for 30 min at 4°C. In all incubation steps cells were protected from light. Fluorescence minus one (FMO) controls were stained alongside samples. Fluorochrome-conjugated mAbs used were anti-mouse CD69-BB700 (H1.2F3, BD Biosciences), CD44-FITC (IM7, Biolegend), CD62L-BV650 (MEL14, BD Biosciences), CD11a-BV605 (2D7, BD Biosciences), KLRG1-PE-Cy7 (2F1, Biolegend), PD-1-BV421 (29F.1A12, Biolegend), CD8-BUV496 (53-6.7, BD Biosciences), CD3-BUV395 (17A2, BD Biosciences), CD4-APC-H7 (GK1.5, BD Biosciences) and CD45-APC (30-F11, Biolegend). Data were acquired with a BD FACSymphony A5 Cell Analyzer in the Flow Cytometry Core at Weill Cornell Medicine and analyzed with FlowJo v10.8 software (BD Life Sciences).

#### Relapse study

B6.Cg-*Prkdc^scid^*/SzJ mice (The Jackson Laboratory, 001913) were aerosol infected with the triple-kill-switch (TKS) strain with 50-100 CFU. The mice were fed with permissive chow (2000 ppm doxycycline and 275 ppm TMP) starting 3 days before infection and through 14 days post. After day 14, mice were switched to regular chow till day 98 to clear infection. The mice were then back on permissive chow for 10 weeks till day 168 to assess relapse. The bacteria burden was assessed at day 1, 14, 21, 28, 56, 98 and 168 from lung and spleen homogenates.

### Macaque studies

Six Mauritian cynomolgus macaques were challenged with 6 CFU Mtb Erdman dual-lysin strain via bronchoscope. Doxycycline was administered daily in food treats beginning 1 day prior to Mtb infection. Doxycycline was discontinued after 2 weeks for 4 macaques while the other two macaques were administered doxycycline for a total of 6 weeks. Comprehensive necropsies were performed at 6 weeks post-challenge. Gross pathology scores reflect the presence of any granulomas or other pathologies, size of lymph nodes, and any other TB related disease. Score of 7 is considered normal/uninfected. Number of involved lymph nodes reflects thoracic lymph nodes with visible granulomas or that are CFU+. Any TB related pathologies (such as granulomas), all lung lobes, all thoracic and peripheral lymph nodes, spleen and liver were harvested and individually processed into single cell suspensions. Samples were plated on aTc containing 7H11 plates, incubated at 37C with 5%CO_2_ and counted at 3 and 6 weeks post-plating.

## Acknowledgements

We thank Michael Chao for discussions during this work. We thank Rodrigo Aguilera Olvera, Heather Kim and Larry Pipkin for technical help with mouse infections. We appreciate the dedication and expertise of the veterinary and research technicians and PET CT imaging personnel at the University of Pittsburgh School of Medicine. This work was supported by NIH R01 AI135629 and Bill & Melinda Gates Foundation INV-009003 and OPP1135516.

## Notes

### Competing Interest Statement

The authors have declared no competing interest.

