## Supplement Figure for "Development of an Engineered *Mycobacterium tuberculosis* Strain for a Safe and Effective Tuberculosis Human Challenge Model"

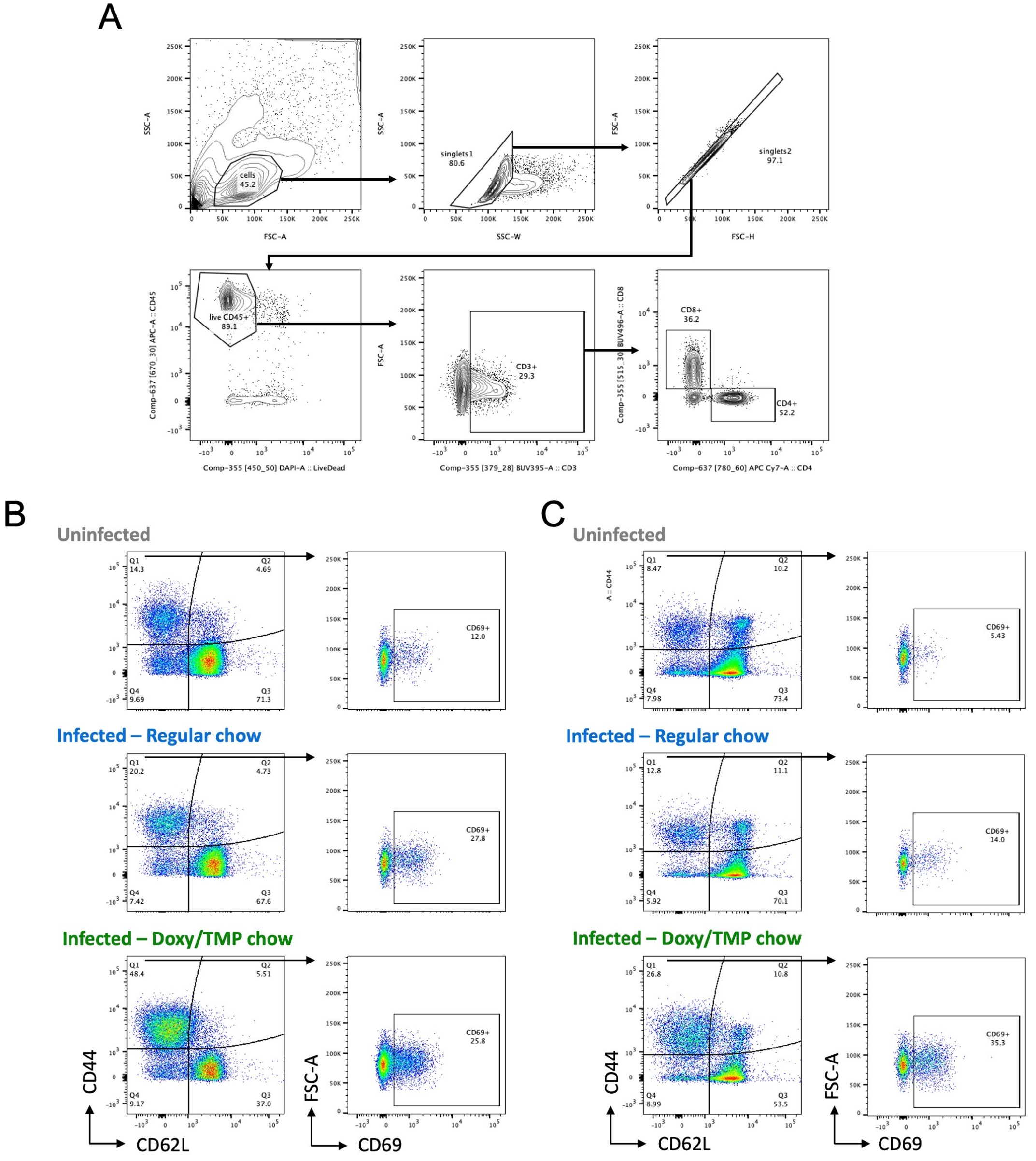


**Supplement figure 1.** Gating strategy for flow cytometry analysis of mouse lung cells. (**A**) General gating strategy for CD4 and CD8 T cells. (**B**) Gating strategy for CD4 T cell memory populations. Central memory = CD62L$^{+}$ CD44$^{+}$ (Q2), effector memory = CD62L- CD44$^{+}$ (Q1), resident memory = CD62L$^{-}$ CD44$^{+}$ CD69$^{+}$. (**C**) Gating strategy for CD8 T cell memory populations. Central memory = CD62L$^{+}$ CD44$^{+}$ (Q2), effector memory = CD62L$^{-}$ CD44$^{+}$ (Q1), resident memory = CD62L$^{-}$ CD44$^{+}$ CD69$^{+}$.

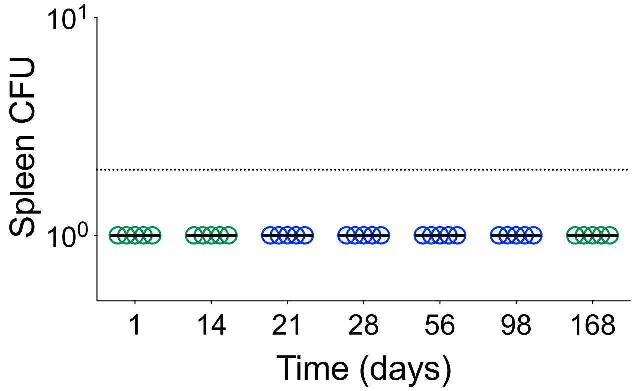


**Supplement figure 2.** Spleen CFU in the SCID experiment. The SCID mice were fed permissive chow for 14 days, followed by restrictive chow till day 98. After that, the mice received permissive chow till day 168. Spleen CFU were enumerated on 7H11 plates.
